# Microbiota control of maternal behavior regulates early postnatal growth of offspring

**DOI:** 10.1101/2020.06.02.130682

**Authors:** Yujung Michelle Lee, Martina Wallace, Jivani M. Gengatharan, Annalee J. Furst, Lars Bode, Christian M. Metallo, Janelle S. Ayres

## Abstract

Maternal-offspring bonding and maturation of maternal behavior is necessary during the early postnatal period of an infant to promote optimal development and growth. The regulation of maternal behavior is multifactorial relying both on sensory cues including auditory and olfactory signals emitted by the infant and received by the mother, as well as proper neuroendocrine responses including that of the oxytocin system. The intestinal microbiota has emerged as a critical regulator of growth and development in the early postnatal period of an individual’s life. While, this is best appreciated in the context of direct interactions between the microbiota and the infant, the microbiota can indirectly influence postnatal growth and development by regulating maternal factors. Here, we describe the identification of an intestinal *E. coli* strain that is pathogenic to the maternal-offspring system during the early postnatal stage of life and results in growth stunting of the offspring. However, rather than having a direct pathogenic effect on the infant, we found that this particular *E. coli* strain was pathogenic to the dams by interfering with the maturation of maternal behavior. The poor maternal behavior resulted in malnourishment of the pups and impaired IGF-1 signaling leading to the consequential stunted growth. Our work provides a new understanding of how the microbiota regulates postnatal growth and an additional variable that must be considered when studying the regulation of maternal behavior.

## Introduction

The early postnatal period of life is a critical stage for growth and development. During this phase there is rapid maturation of all physiologies including immune, metabolic, endocrine and neural processes, the proper function of which is necessary for healthy growth and development. Maturation of these pathways is dependent on internal factors including genetic, physiological and epigenetic mechanisms as well as external environmental cues. Disruption of any of these internal or external factors can lead to states of nutritional challenge, where the infant is undernourished leading to the adverse development of these processes, resulting in stunted growth or wasting. The mammalian microbiota – the trillions of microbes that reside on the body surfaces exposed to the environment – has emerged as a critical regulator of postnatal growth and development, influencing both wasting and growth stunting pathogenesis. In a mouse model of infection induced muscle wasting, a microbiota *E. coli* was found to be protective against muscle wasting via manipulation of an innate immune – endocrine signaling axis that enables the skeletal muscle to maintain a healthy mass during the infection (Schieber et al., 2015). In children, malnutrition was associated with an immature fecal microbiota profile (Subramanian et al., 2014) and colonization of germ free mice with the fecal microbiota of undernourished children was sufficient to cause growth stunting in the gnotobiotic mice (M. I. Smith et al., 2013). Conversely, gavage of gnotobiotic mice with malnourished microbiota with microbiota constituents harvested from healthy donors was sufficient to promote weight and lean mass gain (Blanton et al., 2016). The mechanisms by which the intestinal microbiota will influence early postnatal development and growth are likely complex and multifactorial, however one common theme that has emerged from these initial studies is that the intestinal microbiota is an important regulator of the growth hormone (GH, somatotropin)/insulin-like growth factor-1 (IGF-1) axis, that is important for promoting healthy growth of the host (Schwarzer et al., 2016; Suzawa, Muhammad, Joseph, & Bland, 2019; Yan et al., 2016).

While the focus has been to understand how dysbiotic microbiotas, and the direct interactions between the microbiota and the infant can cause impaired growth and development, maternal factors that are important for healthy postnatal early development and growth will also be regulated by the microbiota. Breast milk contains a microbiota that likely is important for the proper nutritional composition of the milk (Hunt et al., 2011). Maternally derived IgA antibodies that are administered to infants during the early postnatal period and important for shaping the early microbiota of the offspring are regulated by the microbiota (Gopalakrishna et al., 2019; Koch et al., 2016; Lindner et al., 2015). A necessary factor that regulates healthy development and growth of offspring during the early postnatal period is maternal-offspring interactions and bonding. These interactions are essential for providing nutrients to and nurturing of the offspring and are largely regulated by the oxytocin neuroendocrine system that plays a critical role in promoting maternal behavior after birth. The oxytocin system and maternal behaviors are regulated by a number of sensory cues including auditory, visual and olfactory cues. It is currently unknown whether the intestinal microbiota controls the maturation of maternal behavior but given the critical role for the microbiota in the regulation of other behaviors, these seems likely. Lines of investigation to establish a role for the intestinal microbiota in regulating maternal behavior and maternal-offspring interactions and the consequences for offspring growth and development are necessary.

Here we describe our analysis that furthers our understanding of the mechanisms by which the microbiota regulates growth and development during the early postnatal stage of life. We describe a candidate *E. coli* screen in gnotobiotic mice and demonstrate that there is significant variability in the ability of *E. coli* strains to regulate linear growth and body weight in a murine postnatal model. We identified one *E. coli* strain, MG1655, that was pathogenic to infant mice, causing stunted linear growth and body weight gain. Consistent with previous reports, we found this stunting to be dependent on impaired IGF-1 signaling in the pups. However, rather than this being due to direct interactions between the microbe on the pup, we found that the defects in endocrine signaling and consequential stunting to be due to impaired maternal factors. Specifically we found that dams monocolonized with *E. coli* MG1655 exhibited poor maternal behavior, resulting in malnourishment of their offspring, which likely contributed to the attenuated IGF-1 signaling in the offspring. Cross-fostering of *E. coli* MG1655 pups onto dams that exhibited optimal maternal behavior was sufficient to rescue the defective IGF-1 and growth stunting phenotypes. Altogether, our findings support a model by which the intestinal microbiota is necessary for the proper maturation of maternal behaviors after birth and that disruption of these behaviors leads to malnourishment of offspring that results in growth stunting.

## Results

### A candidate screen to identify *E. coli* strains that differentially affect postnatal growth

*Escherichia* species are one of the first colonizers of the infant gut after birth (Houghteling & Walker, 2015). The pan-genome of *Escherichia colis* comprises ~16,000 genes of which represent the core *E. coli* genome (Lukjancenko, Wassenaar, & Ussery, 2010). Thus 90% of the pan-genome varies amongst *E. coli* strains. We hypothesized that this genomic variation can lead to phenotypic variation in various aspects of postnatal health including growth and development. To test this, we performed a candidate microbiome *E. coli* screen in gnotobiotic mice. We gavaged adult female germ-free Swiss Webster mice with one of four different human or mouse-derived *E. coli* strains to establish monocolonized gnotobiotic mice (**Figure 1A and B**). We then bred these mice to male Swiss Webster mice that were monocolonized with the respective *E. coli* strain. F1 progeny were then bred and the resulting F2 progeny were screened for body weight and longitudinal growth at postnatal day (PND) 21 and compared to germ free and specific pathogen free (SPF) Swiss Webster control mice of the same age and litter size (**Figure 1C-F**). SPF mice had greater longitudinal growth compared to germ free mice, indicating that the microbiota promotes linear growth during the postnatal phase (**Figure 1C and D**). Similarly, SPF mice exhibited significantly greater body weight compared to germ free mice indicating that the microbiota is necessary for normal growth during the postnatal stage as has been previously described (Schwarzer et al., 2016; Storelli et al., 2011; Subramanian et al., 2014).

**Figure 1:**
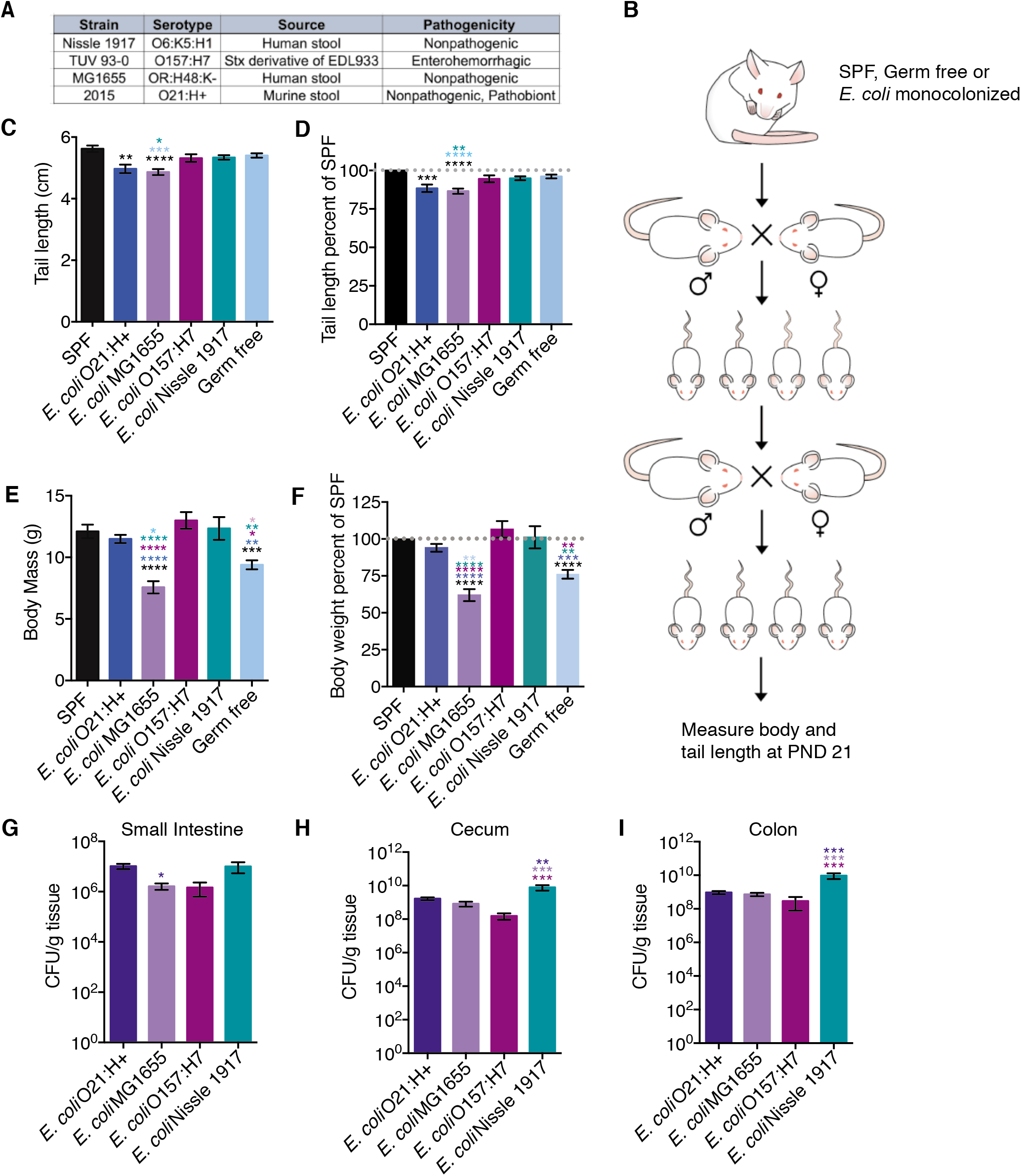
A candidate screen to identify *E. coli* strains that differentially affect postnatal growth. (A) Details of *E. coli* strains used for the candidate screen. (B) Experimental scheme of the *E. coli* candidate screen. Each group of germ free mice were orally gavaged with single strain culture of *E. coli* (5×10^8^ CFU per mouse), mated, and maintained in gnotobiotic isolators. The monocolonized 6 to 8-week-old F1 progenies were mated and the resulting F2 progenies were analyzed for postnatal development. (C) Tail length measurements taken at P21 to assess linear growth of gnotobiotic litters. SPF (n = 22), O21:H+ (n = 7), MG1655 (n = 27), O157:H7 (n = 6), Nissle 1917 (n = 10), and GF (n = 25). (D) tail length measurements shown in (C) as percentage of average SPF length. (E) Body mass of E. coli gnotobiotic litters at P21. SPF (n=27), O21:H+ (n=45), MG1655 (n=32), O157:H7 (n=6), Nissle 1917 (n=10), and GF (n=40). (F) body weight in (E) as percentage of average SPF weight. (G-I) Intestinal colonization levels of bacteria at P21 (n = 10-18 per condition). Statistical significance was determined using one-way ANOVA with post Tukey test for pairwise analyses. *p<0.05, **p<0.01, ***p<0.001, ****p<0.0001. Error bars represent mean +/- SEM. *E. coli* O21:H+ described in (Ayres, Trinidad, & Vance, 2012; Schieber et al., 2015); *E. coli* MG1655 described in (Anderson, 1975); *E. coli* O157:H7 described in (Ferdous et al., 2015) and *E. coli* Nissle described in (Reister et al., 2014).

From our screen, we found that all *E. coli* strains were capable of stably colonizing the gastrointestinal tract (**Figure 1G, H, and I**). Colonization with any of the tested *E. coli* strains was not sufficient to promote linear growth of pups. Mice monocolonized with *E. coli* O157:H7 or Nissle exhibited comparable linear growth to germ free mice (**Figure 1C** and **D**). Mice monocolonized with *E. coli* O21:H+ or MG1655 exhibited reduced linear growth compared to germ free pups (**Figure 1C** and **D**). *E. coli* MG1655 had the most dramatic negative effect on linear growth with pups exhibiting 10% reduction in linear growth compared to germ free mice and ~14% reduction compared to SPF mice (**Figure 1D**). This indicates that monocolonization with any of the *E. coli* strains tested was not sufficient to promote linear growth observed in SPF mice, and that some strains can be pathogenic to linear growth.

Interestingly, we found that the effects the *E. coli* strains had on postnatal linear growth could be decoupled for their effects on weight gain. Pups monocolonized with *E. coli* Nissle, *E. coli* O21:H+ or *E. coli* O157:H7 had significantly higher body weight compared to germ free mice and comparable weights to SPF mice indicating that colonization with one of these strains was sufficient to promote normal body weight gain (**Figure 1E and F**). By contrast, colonization with the strain *E. coli* MG1655, resulted in a reduced weight phenotype compared to germ free mice, indicating that colonization with *E. coli* MG1655 had a pathogenic effect on both postnatal weight and linear development. *E. coli* MG1655 pups exhibited body weights that were ~15% less than germ free pups and ~40% less than SPF pups of the same age and from similar litter size. Taken together, our results demonstrate that intestinal *E. coli* strains have differential effects on postnatal growth in the mouse and fall into two classes: 1) strains that have no effect on linear growth and positive effects on weight gain; 2) strains that have a pathogenic effect on linear growth but are sufficient to promote normal weight gain, indicating that linear growth can be decoupled from weight; and 3) strains that have pathogenic effects on both linear growth and weight gain.

### Postnatal care provided by *E. coli* MG1655 colonized dams results in growth stunting

From our screen, we determined that pups birthed by *E. coli* MG1655-monocolonized dams exhibited a growth stunting phenotype (**Figure 1C-F**). *E. coli* MG1655 is a derivative of the human isolate K-12 strain that was isolated from the stool of a diphtheria patient in the 1920s (Anderson, 1975). Since its isolation, it has been widely utilized in laboratories as a standard, non-pathogenic *E. coli* strain (Blattner et al., 1997; Hayashi et al., 2006; H. W. Smith, 1975). Thus, we were surprised to reveal a pathogenic effect of *E. coli* MG1655 for mouse development in this maternal-offspring system. Low birth weight is a risk factor for growth stunting (Kelleher et al., 1993). We found that *E. coli* MG1655 pups had comparable birth weights to both germ free and *E. coli* O21:H+ pups, indicating that the failure to thrive phenotype observed in *E. coli* MG1655 pups is not due to low birth weight (**Figure 2A**), We tracked the rate of weight gain of pups for 7-8 weeks post birth and found that from P0-P21 when pups are with their mothers, *E. coli* MG1655 pups exhibited a reduced growth rate compared to SPF, *E. coli* O21:H+ and germ free pups. After weaning at P21, the growth rate of *E. coli* MG1655 mice increased, with female mice reaching comparable weights as control mice around 5 weeks old and males reaching comparable weights as control mice around 8 weeks old (**Figure 2B** and **C**).

**Figure 2:**
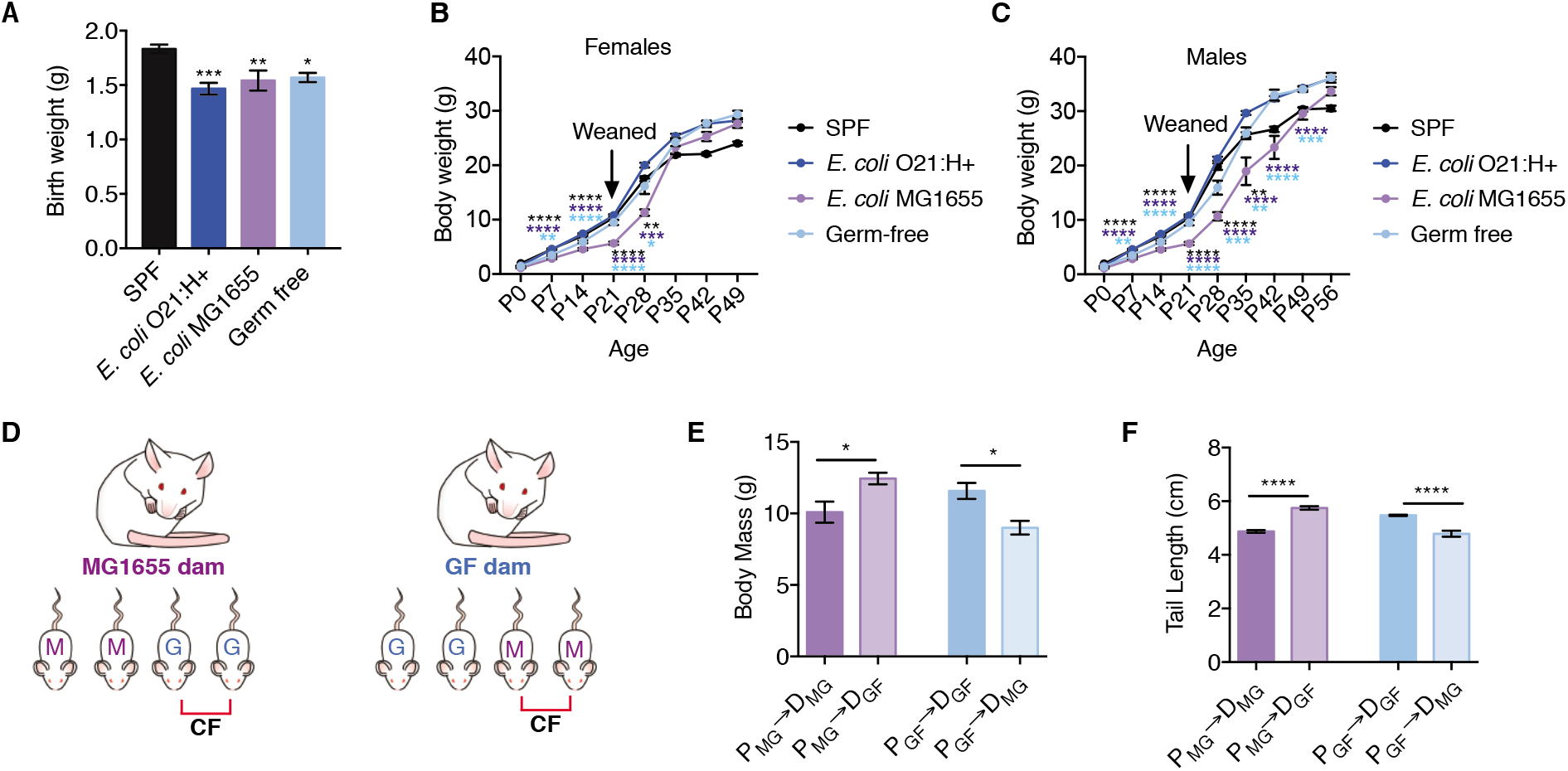
Cross fostering rescues stunted growth exhibited by *E. coli* MG1655 pups. (A) Birth weights of pups (n=10-12 pups per condition). (B-C) Body weights at birth (P0) to adulthood (P49-P56) for (B) females (n=4-11 mice per condition per timepoint) and (C) males (n=5-11 mice per condition per timepoint). From P0-P21, the data represent the weights of both male and female littermates and therefore show the same data in panel B and panel C for those time points. Males and females were separated at the time of weaning and data points beginning at P28 represent the weights of the sex indicated. (D) Experimental scheme of maternal cross-fostering. Immediately after birth, half of the litter is fostered by a different dam (MG or GF) and half of the litter remains with their biological dam. (E) body weight of pups at P21 after cross-fostering (P = pup, D = dam, n = 4 per condition). (F) tail length of cross-fostered pups at P21, (n = 4 per condition). Statistical significance was determined using one-way ANOVA with post Tukey test for pairwise analyses, *p<0.05, **p<0.01, ***p<0.001, ****p<0.0001. Error bars represent mean +/- SEM.

Growth during the pre-weaning phase in a maternal-offspring system is dependent on both maternal and offspring factors. Thus, the pathogenic effects of *E. coli* MG1655 could be due to direct effects of the microbe on the pups or indirectly by influencing maternal factors that are important for offspring growth. To distinguish between these two possibilities, we performed cross fostering experiments where we placed *E. coli* MG1655 pups with germ free dams and germ free pups with *E. coli* MG1655 dams and monitored pup growth (**Figure 2D**). Remarkably, we found cross fostering of *E. coli* MG1655 pups on germ free dams was sufficient to prevent the growth stunting phenotype in these pups. *E. coli* MG1655 pups fostered by germ free dams were comparable in weight and linear length to germ free pups cared for by the same germ free dams and ~120% the weight and length (tail length) of *E. coli* MG1655 pups cared for by *E. coli* MG1655 dams (**Figure 2E** and **F**). Germ free pups fostered by *E. coli* MG1655 dams exhibited a stunted phenotype comparable to *E. coli* MG1655 pups cared for by *E. coli* MG1655 dams and were 80% of the average body weight and tail length of germ free pups with germ free dams (**Figure 2E** and **F**). Thus, the growth stunting phenotype in *E. coli* MG1655 pups was due to indirect effects of *E. coli* MG1655 that disrupt maternal factors that are important for offspring growth.

### *E. coli* MG1655 pups are malnourished and exhibit reduced milk consumption

Growth stunting is caused by a lack of nutrients. Low muscle and fat mass are important indicators of malnutrition (Fischer, JeVenn, & Hipskind, 2015). Consistent with this, body composition analysis revealed that *E. coli* MG1655 pups had significantly reduced fat and muscle composition compared to SPF, germ free and *E. coli* O21:H+ pups (**Figure 3A and B**). This suggests that *E. coli* MG1655 negatively affected offspring growth by disrupting maternal derived nutrients. This can be due to changes in the quality or quantity of milk provided by dams to their pups. Sialylated milk oligosaccharides promote microbiota-dependent growth in infants (Charbonneau et al., 2016). To determine whether milk produced by *E. coli* MG1655 dams had altered levels of milk oligosaccharides, we quantified sialyl(α2,3)lactose (3SL) and sialyl(α2,6)lactose (6SL) which are the only milk oligosaccharides in mice (Fuhrer et al., 2010; Weiss & Hennet, 2012). We found no differences in 3SL or 6SL levels in milk collected from SPF, germ free, and *E. coli* MG1655-monocolonized dams (**Figure 3C and D**), suggesting that *E. coli* MG1655 does not cause growth stunting in pups by negatively affecting the ability of dams to produce milk composed of the major sialylated milk oligosaccharides.

**Figure 3:**
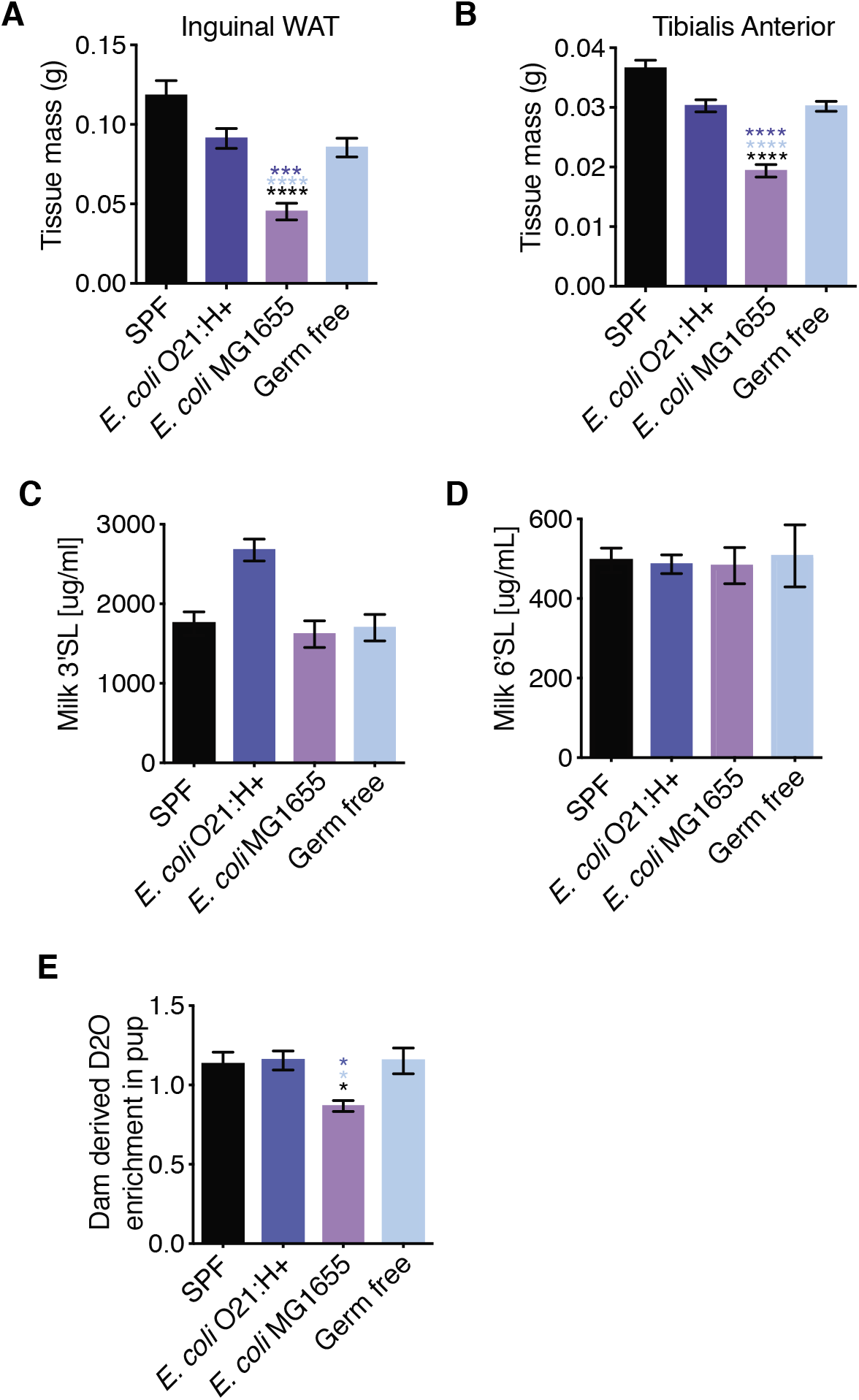
E. coli MG1655 pups are malnourished and exhibit reduced milk consumption. (A) Inguinal white adipose tissue and (B) skeletal leg muscle tibialis anterior (TA) masses of P21 pups, (n = 10-31 mice per condition). (C-D) Levels of (C) 3’SL and (D) 6’SL in milk collected from dams with litters aged P9-P10 (during peak milk production), (n = 4-6 per condition). (E) Percent enrichment of D_2_O detected in pup sera after 48hrs of nursing on D_2_O treated dams (n=19-24 per condition). Statistical significance was determined using one-way ANOVA with post Tukey test, *p<0.05, **p<0.01, ***p<0.001, ****p<0.0001. Error bars represent mean +/- SEM.

We next determined if the growth stunting phenotype exhibited by *E. coli* MG1655 pups was associated with overall reduced feeding. To do this, we utilized a D_2_O-based labeling approach to quantify milk consumption of pups. Lactating dams were injected with a bolus of D_2_O-saline and then maintained on 8% D_2_O water for stable enrichment of D_2_O in the body. Pups were allowed to nurse on D_2_O-enriched dams for 48 hrs, after which we quantified the percent enrichment of D_2_O in the serum of individual pups as a proxy for milk consumption amount over the 48-hour testing period. We found that D_2_O enrichment in the sera of *E. coli* MG1655 pups was significantly reduced, compared to SPF, germ free, and gnotobiotic *E. coli* O21:H+ mice (**Figure 3E**), suggesting that in this *E. coli* MG1655 dam-offspring system, growth stunting is associated with disturbances in the quantity of milk provided by the dam to offspring.

### *E. coli* MG1655 dams exhibit impaired maternal behavior

There are physiological and behavioral maternal factors that influence milk availability for offspring. We performed a nipple function and suckling assay to determine if *E. coli* MG1655 monocolonized dams had defects in milk production or nipple function (Al Ain, Belin, Schaal, & Patris, 2013). Pups were removed from their mothers to fast, after which they were weighed and returned to their sedated mothers that were turned on their backs with nipples exposed. The pups were placed next to their mothers and allowed to latch on her nipples and suckle. The pups were allowed to feed for one hour, after which they were weighed. The change in weight was used as an indirect measurement for the amount of milk consumed. We found that the change in weight exhibited by *E. coli* MG1655 pups was comparable to SPF and *E. coli* O21:H+, all of which were greater than the weight change exhibited by germ free pups (**Figure 4A**). This suggests that *E. coli* MG1655 dams have functional nipples and produce and secrete comparable volumes of milk in a defined feeding period. It also demonstrates, consistent with our cross fostering assay, that the reduced milk intake exhibited by *E. coli* MG1655 pups is not due to defects in the pups ability to latch and suckle.

**Figure 4:**
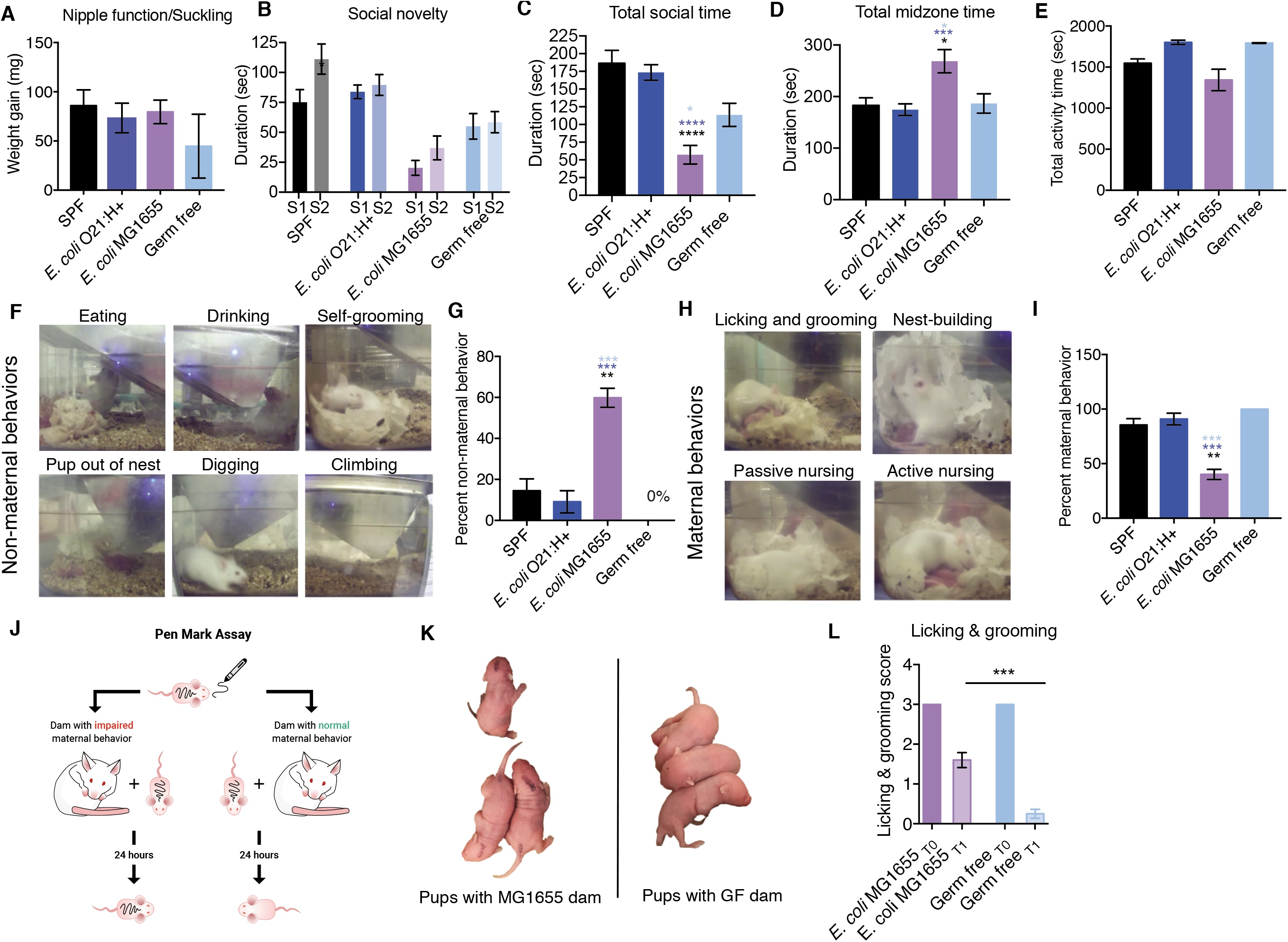
Impaired maternal behavior in MG1655 dams. (A) Weight gain of pups after feeding on sedated dams for one hour (n = 3-8 per condition). (B) Social novelty test (n=13-30 per condition). S1 = familiar mouse and S2 = novel mouse. (C) Total socialization time calculated from social novelty test in B (n=13-30 per condition). (D) Total time in midzone calculated from the social novelty test in B (13-30 per condition). (E) Dams were recorded for 30 minutes and the total amount of active time was measured (n=2-3 per condition). (F) Representative images of non-maternal behaviors. (G) Dams were recorded and the total amount of time performing the non-maternal behaviors in (G) was measured. (H) Representative images of maternal behaviors. (I) Dams were recorded and the total amount of time performing the maternal behaviors in (H) was measured. (J) Schematic demonstrating the “pen mark” assay for quantification of licking and grooming. (K) Representative images of pups from the “pen mark” assay. Pups cared for by E. coli MG1655 dams have visible pen mark 24 hrs later while pups with germ free dams have barely visible pen mark. (L) Quantification of the pen mark assay (n=5-6 pups per condition). Statistical significance was determined using one-way ANOVA with post Tukey test or unpaired student’s t-test for pairwise analyses, *p<0.05, **p<0.01, ***p<0.001, ****p<0.0001. Error bars represent mean +/- SEM.

We next considered the possibility that *E. coli* MG1655 had behavioral defects that influenced their ability to provide adequate care and nutrients to their offspring. We first examined the social behavior of *E. coli* MG1655 monocolonized adult females and control mice. We performed a three-chamber social novelty test to assess social behavior. Mice prefer to spend more time with another mouse and prefer a novel social subject over a familiar one (social novelty). In this test, the test subject is allowed to freely explore inside a three-chambered box with openings between the chambers. One chamber houses a familiar mouse and another chamber houses a novel mouse. The amount of time the test mouse interacts with the familiar and novel mouse are recorded. We found that while *E. coli* MG1655-monocolonized mice preferred to spend time with the novel mouse, the total amount of time spent socializing by *E. coli* MG1655 was significantly reduced compared to SPF, *E. coli* O21:H+ monocolonized and germ free mice (**Figure 4B** and **C**). Instead *E. coli* MG1655 mice spent significantly more time not socializing and contained within the midzone of the assay apparatus compared to the three control conditions (**Figure 4D**). From video recording analysis, we determined that the decreased socialization time and increased midzone time was not due to reduced activity or mobility issues, as *E. coli* MG1655 monocolonized dams exhibit comparable activity times as SPF dams (**Figure 4E**).

We next considered that *E. coli* MG1655 dams had impaired maternal behavior. We video recorded dams with their litters in 30 minute segments of time from P2-P4 and measured the amount of time the dams spent performing non-maternal and maternal behaviors (**Figure 4F-I**). For non-maternal behaviors, we quantified the amount of time the dam spends digging and climbing, eating and drinking, self-grooming, and when pups were found outside of their nest (**Figure 4F**) (Capone, Bonsignore, & Cirulli, 2005). For maternal behaviors, we looked for nest building, licking and grooming, passive nursing, and active nursing (**Figure 4H**). We found that *E. coli* MG1655 dams exhibited greater time performing non-maternal behaviors compared to our three control conditions and significantly less time performing maternal behaviors compared to SPF, *E. coli* O21:H+ and germ free dams (**Figure 4F-I**). We next developed the “pen mark” assay that could allow us to better quantify maternal behaviors such as licking and grooming (**Figure 4J**). Briefly, pups were marked on two distinct anatomical regions (head and rear) using a VWR marker and placed back in their home cages containing their respective dams. After 24 hours, each pup was scored for the intensity of the marks on each region of the body as a way to measure the dam’s licking and grooming behavior (**Figure 4K** and **L**). We found that pups that were with *E. coli* MG1655 dams exhibited greater pen mark intensity after 24hrs compared to pups that were placed with germ free dams indicating that *E. coli* MG1655 dams had reduced licking and grooming behaviors in the 24 hr period compared to germ free dams (**Figure 4K** and **L**). In conclusion, the growth stunting and impaired milk consumption exhibited by *E. coli* MG1655 pups is associated with impaired maternal behavior in *E. coli* MG1655 dams.

### *E. coli* MG1655 pups have attenuated IGF-1 signaling

The growth hormone (GH) – liver IGF-1 axis is a critical regulator of postnatal growth and is regulated by the nutritional state of the organism (Hintz, Suskind, Amatayakul, Thanangkul, & Olson, 1978; Stratikopoulos, Szabolcs, Dragatsis, Klinakis, & Efstratiadis, 2008). During malnutrition and fasting states, levels of IGF-1 and signaling are reduced (Hawkes & Grimberg, 2015; Moore et al., 2008). We hypothesized that the failure to thrive and malnourished phenotype of *E. coli* MG1655 pups was associated with reduced IGF-1 activity. To determine this, we first looked at IGF-1 signaling in the liver and found that there was reduced phosphorylation of the serine/threonine kinase called Akt in *E. coli* MG1655 pups indicating that these pups had reduced IGF-1 signaling (**Figure 5A**). Reduced IGF-1 signaling in the *E. coli* MG1655 pups could be due to reduced production of IGF-1 by the liver, reduced secretion and/or stability of IGF-1 in the circulation, or IGF-1 resistance. To distinguish between these possibilities, we first measured the levels of IGF-1 in the livers. We found that *E. coli* MG1655 pups had comparable levels of IGF-1 in their livers (**Figure 5B**). Furthermore, the levels of IGF-1 in the white adipose tissue, a second tissue source for IGF-1 that can regulate body size (Kloting et al., 2008), were not significantly different in *E. coli* MG1655 pups compared to the three control conditions (**Figure 5C**). By contrast, serum levels of IGF-1 were significantly reduced in *E. coli* MG1655 pups compared to SPF, *E. coli* O21:H+ and germ free pups, suggesting that the reduced IGF-1 signaling in *E. coli* MG1655 pups is due to impaired secretion and/or stability of circulating IGF-1 (**Figure 5D**).

**Figure 5:**
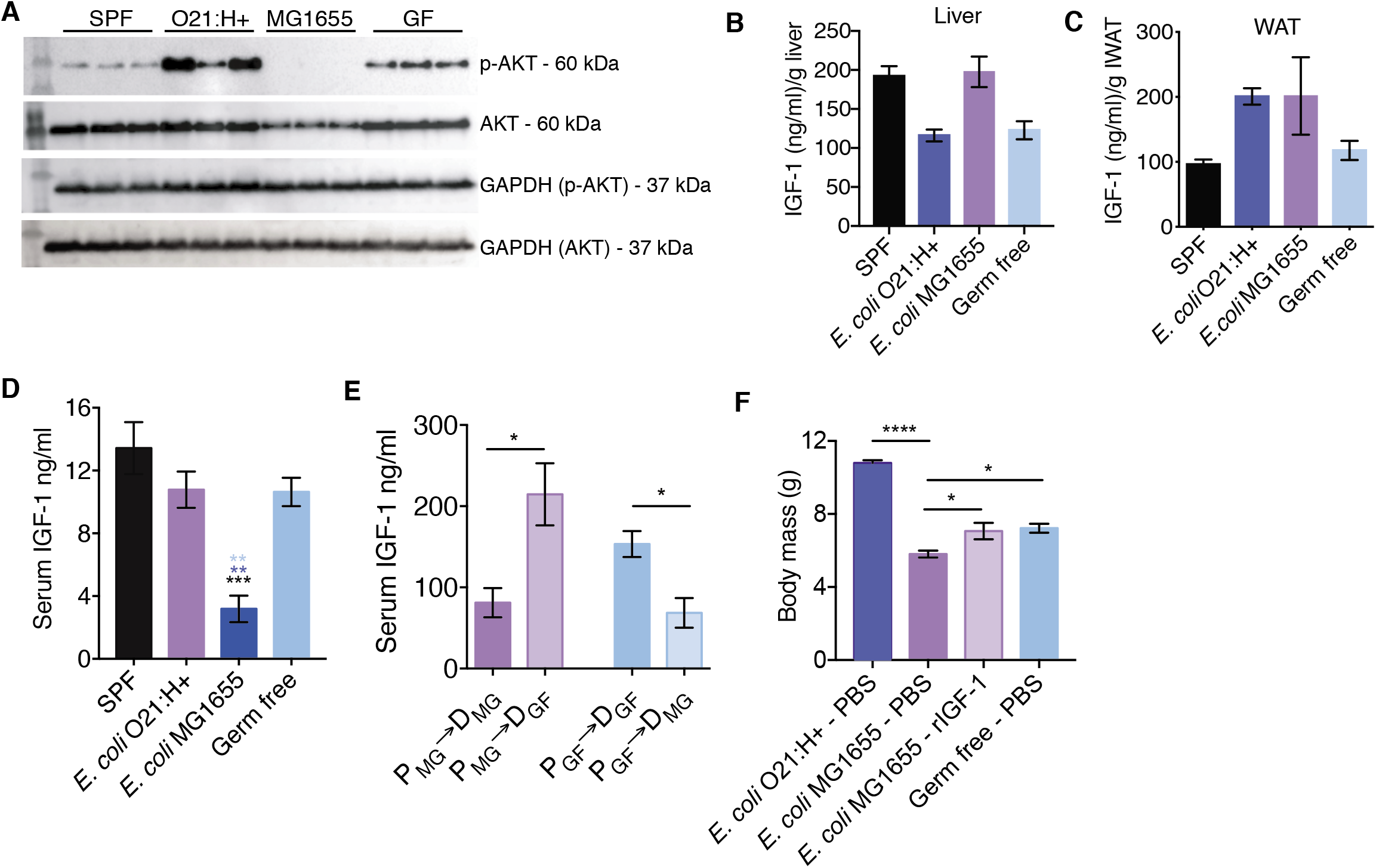
Impaired IGF-1 signaling in MG1655 neonates. (A) IGF-1 signaling levels in liver at P3 determined by measuring phosphorylated AKT and AKT levels. GAPDH for loading control (n = 3 per condition). (B) IGF-1 protein levels in liver at P3 (n = 6-23 per condition). Serum IGF-1 levels, n = 8-16 per condition. (C) Adipose tissue IGF-1 protein levels at P3, n = 6-11 per condition (D) Serum IGF-1 levels at P3 (n = 4-7 per condition). (E) Serum IGF-1 levels at P21 from cross-fostering experiments (n=4 per condition). (F) Body weight measurements of animals treated with daily intraperitoneal injection of mouse recombinant IGF-1 or PBS vehicle control for 14 days, n = 6-13 per condition. Statistical significance was determined using one-way ANOVA with post Tukey test or unpaired t-test, *p<0.05, **p<0.01, ***p<0.0005, ****p<0.0001. Error bars represent mean +/- SEM.

Our data thus far indicate that the growth stunting phenotype exhibited by *E. coli* MG1655 is associated with reduced IGF-1 levels and downstream signaling. Our data further show that the failure to thrive phenotype is regulated by maternal factors. To determine if the effects of *E. coli* MG1655 on IGF-1 in pups is due direct actions of the microbe on the pups or indirectly via interactions between the microbe and the dam, we measured the circulating levels of IGF-1 in pups from our cross-fostering experiment (**Figure 2D-F**). Cross fostering of *E. coli* MG1655 pups by germ free dams exhibited significantly higher circulating levels of IGF-1 compared to their littermates that were left to nurse on their *E. coli* MG1655 dam (**Figure 5E**). Germ free pups that were fostered by *E. coli* MG1655 dams exhibited significantly reduced IGF-1 serum levels compared to their littermates that were left to nurse on their germ free dam (**Figure 5E**). Thus, monocolonization with *E. coli* MG1655 impairs maternal factors (presumably maternal behavior and the ability of the dam to provide nutrients) of the dam that are necessary for promoting circulating IGF-1 levels in their offspring.

To determine whether exogenous supplementation of IGF-1 could rescue the failure to thrive phenotype in *E. coli* MG1655 pups, we intraperitoneally injected P3 neonates with recombinant IGF-1 (rIGF-1) for 14 days and compared their body weights to those of PBS-injected control groups. We found that *E. coli* MG1655 pups that were injected with exogenous of rIGF-1 had increased growth compared to *E. coli* MG1655 pups that were injected with PBS (**Figure 5F**). The body weight of *E. coli* MG1655 pups treated with rIGF-1 was comparable to that of germ free pups injected with PBS (**Figure 5F**). However, the body weight of rIGF-1-treated pups was significantly reduced compared to *E. coli* O21:H+ pups injected with PBS (**Figure 5F**). Taken together, our data suggest that administration of exogenous IGF-1 was sufficient to rescue *E. coli* MG1655 pups from the stunted growth phenotype enabling them to reach comparable size as germ free mice. However, IGF-1 administration was not sufficient to promote growth comparable to *E. coli* O21:H+ or SPF in *E. coli* MG1655 pups indicating that there are additional microbial factors that are necessary to promote optimal weight gain and that are lacking in the *E. coli* MG1655 maternal offspring system.

## Discussion

We present our approach to investigate the effects of different *E. coli* strains on postnatal development. Our work supports a model by which *E. coli* MG1655 monocolonization impairs the maturation of maternal behaviors in dams after birth. This results in malnourishment of the offspring and negatively affects IGF-1 levels and signaling in the offspring, presumably as a consequence of the malnutrition. Administration of exogenous IGF-1 was sufficient to rescue the pathogenic effects of *E. coli* MG1655 but was not sufficient to promote healthy growth similar to SPF mice and like the other *E. coli* strains tested (**Figure 6**). Our study adds to the ever growing evidence that the microbiota is necessary for optimal growth and development and that it is necessary for normal behavioral processes.

**Figure 6.**
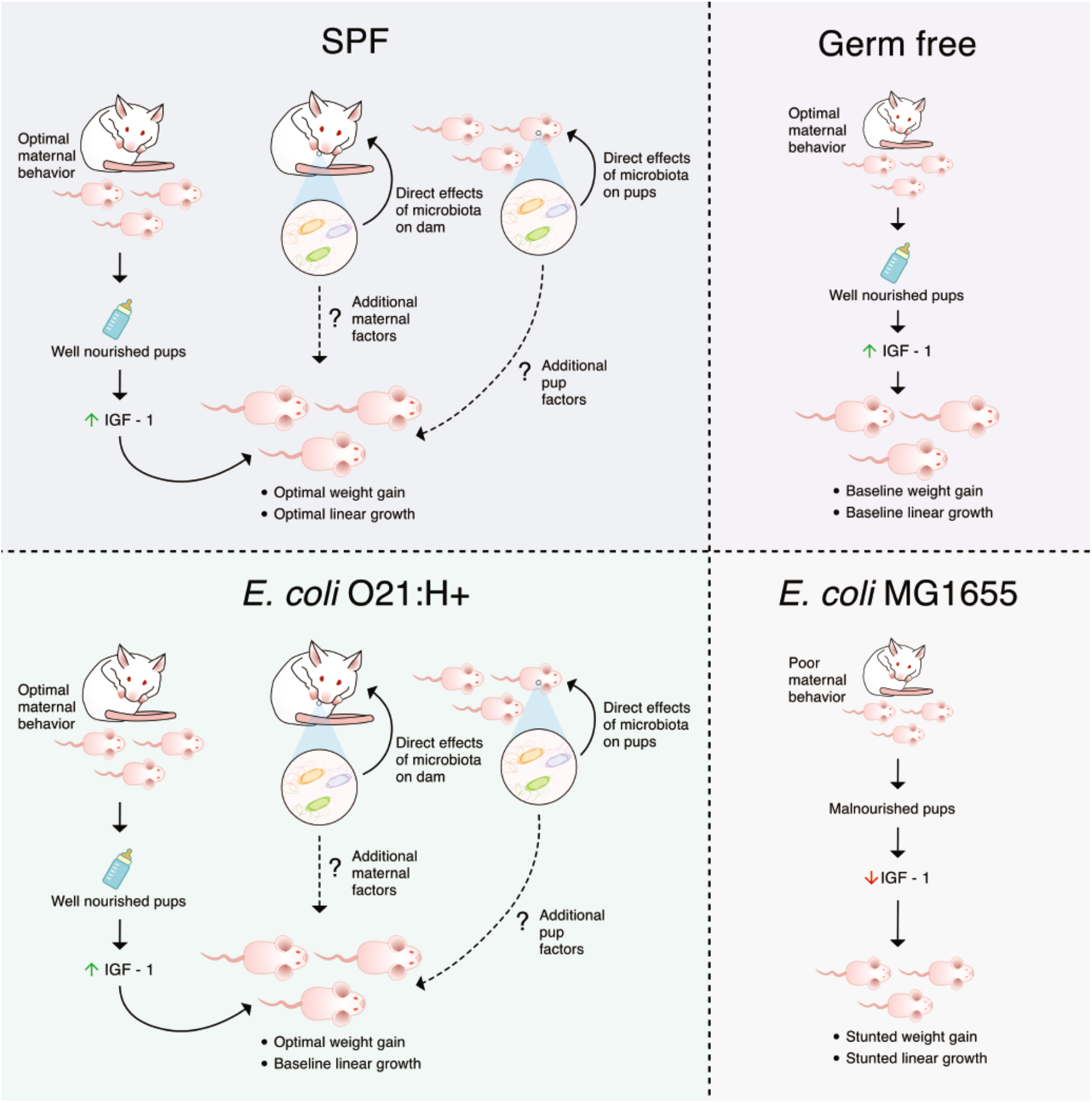
E. coli MG1655 impairs maternal behavior resulting in offspring malnutrition and growth stunting. Our work supports a model by which *E. coli* MG1655 monocolonization impairs the maturation of maternal behaviors in dams after birth. This results in malnourishment of the offspring and negatively affects IGF-1 levels and signaling in the offspring, presumably as a consequence of the malnutrition. Administration of exogenous IGF-1 was sufficient to rescue the pathogenic effects of *E. coli* MG1655 but was not sufficient to promote healthy growth similar to SPF mice and like the other *E. coli* strains tested.

Shortly after birth, the infant intestine becomes colonized by maternal and environmentally derived facultative anaerobic bacteria such as *E. coli* that play a critical role in postnatal development. From our candidate *E. coli* screen, we identified strains that varied in their ability to promote normal postnatal growth in an infant mouse model. We found that while none of the *E. coli* strains screened could promote normal linear growth compared to SPF mice, the *E. coli* strains varied in their ability to promote normal weight gain during the pre-weaning period. Colonization with *E. coli* O21:H+, Nissle or O157:H7 was sufficient to promote weight gain comparable to that exhibited by SPF pups and greater than germ free pups. Our data demonstrate that the effects of different microbiota *E. coli* strains have on linear growth and weight gain can be decoupled. Impaired linear growth with normal weight gain in humans is associated with metabolic and cognitive disorders later in life (De Lucia Rolfe et al., 2018; Xie et al., 2019). Furthermore, disruptions in microbiota homeostasis contributes to metabolic and cognitive disorders (Buffington et al., 2016; Kim et al., 2017; Ley et al., 2005). Our findings suggest that one way in which infant intestinal dysbiosis may contribute to health problems later in life may be due to the disruption in the relationship between linear growth and weight gain and adds to the ever growing evidence that the influence of the intestinal microbiota early in life has on multiple aspects of health at later life stages. In adults, numerous disease settings are associated with intestinal dysbiosis characterized by an overrepresentation of *E. coli* and other members of the *Enterobacteriaceae* family, suggesting these bacteria may cause or exacerbate disease states in adults (Baumgart et al., 2007; Shin, Whon, & Bae, 2015). Our findings that *E. coli* O21:H+, Nissle and O157:H7 promote healthy infant postnatal growth suggests that the effects microbes have on the health of their hosts are complex and can vary over the different life stages of the individual – being beneficial for some life stages but pathogenic at other life stages.

We identified one *E. coli* from our screen, *E. coli* MG1655, that was pathogenic to infant mice and resulted in a growth stunted phenotype. In addition to impaired linear growth, *E. coli* MG1655 pups exhibited significantly reduced weight compared to not only SPF mice but also germ free control mice. We found that the growth stunting of *E. coli* MG1655 monocolonized pups was associated with reduced circulating levels of IGF-1 and signaling. Administration of exogenous IGF-1 was sufficient to rescue the pathogenic effects of *E. coli* MG1655 and the resulting stunted phenotype, enabling the *E. coli* MG1655 pups to reach similar body weight of germ free pups. However, *E. coli* MG1655 pups treated with rIGF-1 still did not reach comparable size of pups monocolonized with *E. coli* O21:H+ or SPF pups, suggesting that *E. coli* MG1655 lacks the ability to promote normal weight gain like other *E. coli* strains we used in our study. As *E. coli* MG1655 is a derivative of the original human isolate K-12 strain that was initially isolated from a diphtheria patient, and is now a lab adapted strain, it is possible that it lost factors that enable it to function similarly to our other *E. coli* strains and promote healthy weight gain during the pre-weaning phase of infant mice. Another interesting possibility is that the lipopolysaccharide (LPS) of MG1655 is phenotypically rough and lacks the O antigen, which is its repeated oligosaccharide unit (Browning et al., 2013). Understanding the microbial factors that contribute to *E. coli* MG1655 pathogenicity and those that enable the other *E. coli* strains to promote healthy growth will be of interest for future studies.

Our discovery that an intestinal microbiota species regulates systemic IGF-1 and body size is in agreement with previous findings from our group and others. We previously reported in a mouse model of infection induced muscle wasting that the intestinal microbiota regulates circulating levels of IGF-1 and body size (Schieber et al., 2015). Subsequent studies in mice and fruit flies showed that intestinal microbiota regulation of the growth hormone (GH, somatotropin)/insulin-like growth factor-1 (IGF-1) axis, is important for promoting healthy growth of the host (Schwarzer et al., 2016; Storelli et al., 2011). In these previous studies, the microbiota was shown to regulate IGF-1 through direct interactions with the host. In the current study, we demonstrated that in maternal-infant systems, the intestinal microbiota can regulate growth and IGF-1 levels in the offspring indirectly, via effects on maternal factors. With our cross fostering experiments, we demonstrated that pups that received maternal care from *E. coli* MG1655 dams had reduced circulating IGF-1 regardless of intestinal microbiota status of the pups. By contrast, fostering of *E. coli* MG1655 pups by germ free dams was sufficient to rescue the impaired IGF-1 phenotype in *E. coli* MG1655 pups. We found that dams monocolonized with *E. coli* MG1655 exhibited impaired social and maternal behavior, and we suggest that these behavioral defects contribute to *E. coli* MG1655’s negative effects on infant IGF-1 levels and growth. Maternal care is necessary to provide infants with nutrients and IGF-1 secretion and stability is dependent on the nutritional status of the host (Hawkes & Grimberg, 2015; Hintz et al., 1978; Stratikopoulos et al., 2008). In agreement with this, we found that *E. coli* MG1655 pups consume less milk when receiving maternal care from *E. coli* MG1655 dams compares to pups that receive maternal care from germ free, SPF or *E. coli* O21:H+ monocolonized dams. Thus *E. coli* MG1655 impairment of maternal behavior likely prevents dams from providing adequate nutrients that regulate IGF-1 levels in infants. In future work, it will be important to investigate other aspects of maternal care including bonding, warmth and grooming, which may also contribute to the IGF-1 deficiency in *E. coli* MG1655 pups.

An outstanding question from our study is what is the mechanism by which *E. coli* MG1655 impairs maternal behavior? The oxytocin neuroendocrine system is a critical regulator of maternal behavior in the postnatal period and for social bonding, and is regulated by the intestinal microbiota. In mice, treatment with *L. reuteri* improved deficits in social behavior and was associated with increased hypothalamic expression of oxytocin (Buffington et al., 2016). While the oxytocin neuroendocrine system is a likely candidate, whether *E. coli* MG1655 colonization in dams negatively effects this system remains to be determined. Whether the maternal behavior deficiencies are due to direct effects of *E. coli* MG1655 colonization on brain health or a consequence of developmental defects in *E. coli* MG1655 mice independent of the growth stunting remains to be determined, however our results with the cross-fostering studies suggest it may be the latter. In our crossfostering studies, the germ free dams become colonized with *E. coli* MG1655 when these pups are introduced to the dam. Despite becoming colonized with this microbe, ex-germ free dams were able to rescue the *E. coli* MG1655 pups from the stunted phenotype and IGF-1 deficiencies. These data suggest that short term colonization of dams during the postnatal period *E. coli* MG1655 is not sufficient to induce the impaired maternal behavior and stunting phenotype in infants.

Although often overlooked, a major risk factor associated with childhood growth failure is deficits in maternal neurological health, which can interfere with maternal behaviors. For example, several epidemiological studies of mother-infant pairs in developing countries with high prevalence of childhood undernutrition revealed that there is high association between childhood growth failure and poor maternal mental health (Rahman, Lovel, Bunn, Iqbal, & Harrington, 2004; Ross et al., 2011; Surkan, Kennedy, Hurley, & Black, 2011). This association most likely exists because maternal depression interferes with maternal interest in offspring and emotional quality of nurturing, which limit the offspring from reaching maximum growth capacity (Girma, Fikadu, & Abdisa, 2019). The factors that mediate such relationship between postnatal growth failure and maternal behavior are not defined by one general mechanism, but most likely driven by multiple environmental parameters that contribute to optimal maternal-offspring interactions. Altogether, our study adds an additional layer of complexity to our understanding of maternal behavior and demonstrates that the maternal microbiota will affect maternal behavior and this likely regulates offspring development and growth.

## Acknowledgements

We thank Daniela Michel for assistance with the germ free and gnotobiotic husbandry, as well as technical assistance with experiments done with *E. coli* Nissle and *E. coli* O157:H7 gnotobiotic mice; Randy Hampton, Steve Hedrick, Victor Nizet, and Manuela Raffatellu, and members of the Ayres laboratory for helpful discussions and guidance. This work was supported by a Crohn’s and Colitis Foundation Senior Research Award and NIH R01 AI114929 both awarded to JSA. Funding from the Lynne and Mason Rosenthal/Leo S. Guthman Foundation was awarded to YML.

## Contributions

JSA conceived the idea, provided technical guidance, analyzed the data, made the figures and wrote the paper. YML performed experiments, wrote the material and methods section, plotted the data and made the figures. MW, JMG and CMM developed the D_2_O milk consumption assay, performed the experiments and analyzed the data. AJF and LB developed the milk oligosaccharide assay, performed the experiments and analyzed the data.

## Materials and Methods

### Bacterial Strains

To prepare bacterial inocula for oral gavaging, *E. coli* O21:H+, *E. coli* MG1655, *E. coli* O157: H7, and *E. coli* Nissle 1917 were grown in LB media at 37°C shaking and stored in 25% glycerol/PBS frozen stocks and used directly for gavaging.

### Animals/Gnotobiotic Mouse Model

Germ free Swiss Webster mice were orally gavaged with 5×10^8^ CFU of *E. coli* O21:H+, *E. coli* MG1655, *E. coli* O157: H7, or *E. coli* Nissle 1917 and maintained in sterile semi-flexible isolators under a 12 hour light-dark cycle and given autoclaved standard chow diet and distilled water *ad libitum*. The status of germ free and gnotobiotic conditions were monitored routinely for bacterial, viral, and fungal contaminations. All animal experiments were approved by our institutional IACUC committee.

### Assessment of Development

Body weights of each neonate in a given litter were monitored using Pesola spring scale with crocodile clamp inside germ-free, gnotobiotic and SPF semi-flexible isolators during the entire duration of the experiment. Linear growth of neonates was quantified by measuring the length of tail and nose-to-anus distance. For body composition analyses, inguinal white adipose tissue and tibialis anterior muscle were dissected and weighed on Mettler Toledo scale and weights were recorded.

### Quantification of bacterial colony-forming units in tissues

Colonization levels of all *E. coli*s were quantified by serially diluting intestinal homogenates and plating on Eosin methylene blue agar (EMB)/ampicillin, vancomycin, neomycin and metroidazole (*E. coli* O21:H+), or EMB (*E. coli* MG1655, *E. coli* O157: H7, and *E. coli* Nissle 1917). Plates were incubated at 37°C overnight. Intestinal tissues were homogenized in 1ml sterile PBS supplemented with 1% Triton-X using a bead beater. Some germ-free or gnotobiotic intestinal tissues were homogenized in 5mls sterile PBS supplemented with 1% Triton-X using a Qiagen tissue shredder.

### Neonatal rIGF-1 injections

For rIGF-1 injection experiments, germ free and gnotobiotic mice (*E. coli* MG1655 and *E. coli* O21:H+) were transferred from gnotobiotic isolators to autoclaved airtight IsoCage Bioexclusion System with positive pressure and HEPA filtration (Tecniplast SpA, Buguggiate, Varese, Italy), which were pre-packaged with Sani-Chips bedding (P.J. Murphy) and standard chow prior to autoclaving. Autoclaved distilled water was provided to the transferred mice. All handling of mice housed in IsoCages were performed under laminar flow hood with sterilization procedures using the disinfectant Clidox-S (1:3:1 formulation). 3-day-old pups were intraperitoneally injected with sterile murine rIGF-1 (R&D systems) or sterile PBS daily at a dose of 200μg/kg until weaning age (21-days-old) was reached using a 100μl gastight glass syringe (Hamilton 7656-01) with 34-gauge needle (Hamilton pt style 4, 12-30°, 0.375” length). Daily body weights were measured using an electronic scale for tracking body weight development.

### ELISA

Serum and tissue homogenates were prepared for IGF-1 ELISA. Tissues were frozen in liquid nitrogen and homogenized in lysis buffer (1% Triton X-100 in PBS) using Bead Mill 24 (Fisher Scientific). Lysates were centrifuged and supernatants were used for ELISAs. IGF-1 was measured using anti-IGF monoclonal antibody Clone #126002 and anti-mouse IGF-1 Biotinylated Antibody (R&D systems).

### AKT western blot

Tissues were ground in LN2 and protein was extracted using Life Technologies tissue extraction reagent II. Proteins of interest were probed using pAKT (Ser 473) (193H12) Rabbit mAB #4058T, total AKT Rabbit Ab #9272S and GAPDH Ab (Cell Signaling Technology).

### Food intake measurements

#### Milk intake/Suckling behavior/Nipple function assessment

Pups at PND3, 7, 11, 14, and 18 were fasted for 7 hours by removing them from the house cage containing the lactating dam and were placed on a heat pad. After collecting the fasted whole body weights of each pup, the pups in each litter were returned to the home cage containing anesthetized lactating dam placed on her side with the body tilted so that the lines of nipples would be exposed to the pups. To ensure that pups don’t lose their way to the nipples, a divider/wall was placed rostral to the pectoral nipples and another placed caudal to the inguinal nipples. After observation of the pups to latch to the nipples, the pups were allowed to suckle for milk for 1 hour and their final body weights were measured. The body weight gain of each pup during 1 hour of suckling was used to estimate the milk intake.

#### D_2_O labeling & serum D_2_O enrichment measurement

48 hours prior to termination, dam was I.P. injected with 0.035 mls/g body weight 0.9% NaCl-D_2_O and drinking water was replaced with 8% D_2_O enriched water. Dam was fasted for 6 hours prior to serum collection. At the time of sacrifice, serum from individual pups in the litter was also collected. The ^2^H labeling of water from samples or standards was determined via deuterium acetone exchange. 5 μls of sample or standard was reacted with 4 μls of 10N NaOH and 4 μls of a 5% (v/v) solution of acetone in acetonitrile for 24 hours. Acetone was extracted by the addition of 600 μl chloroform and 0.5 g Na2SO4 followed by vigorous mixing. 100 μls of the chloroform was then transferred to a GCMS vial. Acetone was measured using an agilent DB-35MS column (30 m 3 0.25mm i.d. 3 0.25 mm, Agilent J&W Scientific) installed in an Agilent 7890A gas chromatograph (GC) interfaced with an Agilent 5975C mass spectrometer (MS) with the following temperature program: 60 °C initial, increase by 20 °C/min to 100 °C, increase by 50 °C/min to 220 °C, and hold for 1 min. The split ratio was 40:1 with a helium flow of 1 ml/min. Acetone eluted at approximately 1.5min. The mass spectrometer was operated in the electron impact mode (70 eV). The mass ions 58 and 59 were integrated and the % M1 (m/z 59) calculated. Known standards were used to generate a standard curve and plasma % enrichment was determined from this. All samples were analyzed in triplicate. In order to calculate the pup’s normalized % D_2_O enrichment, the dam’s % D_2_O enrichment and the pup’s weight were taken into account using the following equation:

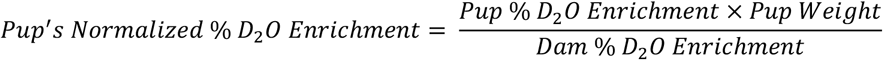

### Social interaction test

Mouse was placed in the central chamber of the three-chamber apparatus and habituated for 5 minutes. Right and left chambers were isolated by using the dividing Plexi glass walls. One control mouse noted as “Social 1” was placed inside a wire containment cup and placed in the left/right chamber and another mouse noted as “Social 2” was placed inside a cup placed in the third chamber. To begin testing, the walls between the chambers were removed and the mouse was allowed to explore each of the three chambers for 10 minutes. For quantification social novelty, ANY-maze software linked to an overhead camera was used to score the amount of the time the mouse spends interacting with the “Social 1” versus “Social 2” subject.

### Licking and grooming assay

To assess maternal behavior of licking and grooming of pups, pups were marked with a lab marker on the top of head and bottom and placed back in their home cage with their respective dams. Photos were taken at T_0_ and T_1_ for records. After 24hrs, the quality of the marks on the pups were assessed based on the following scores:

0 – *no mark visible*
1 – *visible mark is extremely faint*
2 – *visible mark is clear but faint*
3 – *visible mark is clear*

Average of two separate scores (one for head and another for bottom) were plotted for each pup.

### Maternal behavior assessment

Dams with their respective litters were undisturbed and recorded in their home cages during postnatal days 2-4 using a pet camera (Petcube). For each dam, 30-minute video was recorded during AM (9am-12pm) and PM (4pm-7pm) for each testing day. Recorded videos were used to individually score maternal and non-maternal behaviors of each dam by measuring the duration the dam performs the following behaviors:

<u>Maternal behaviors</u>

- Nest-building
- Licking and grooming
- Active nursing
- Passive nursing

<u>Non-maternal behaviors</u>

- Eating and drinking
- Self-grooming
- Digging and climbing
- Pups out of nest

### Statistical analysis

Data are represented as mean SEM. Statistical tests were performed using Graphpad Prism version 6.0. unpaired t test or one way ANOVA with post Tukey test were used for pairwise analyses. Sample size and p values are recorded in each figure legend and p<0.05 was used as a cutoff for determining significance.

## References

Al Ain, S., Belin, L., Schaal, B., & Patris, B. (2013). How does a newly born mouse get to the nipple? Odor substrates eliciting first nipple grasping and sucking responses. Dev Psychobiol, 55(8), 888–901. doi:10.1002/dev.21082 23037148

Anderson, E. S. (1975). Viability of, and transfer of a plasmid from, E. coli K12 in human intestine. Nature, 255(5508), 502–504. doi:10.1038/255502a0 1094298

Ayres, J. S., Trinidad, N. J., & Vance, R. E. (2012). Lethal inflammasome activation by a multidrug-resistant pathobiont upon antibiotic disruption of the microbiota. Nat Med, 18(5), 799–806. doi:10.1038/nm.2729 22522562

Baumgart, M., Dogan, B., Rishniw, M., Weitzman, G., Bosworth, B., Yantiss, R., … Simpson, K. W. (2007). Culture independent analysis of ileal mucosa reveals a selective increase in invasive Escherichia coli of novel phylogeny relative to depletion of Clostridiales in Crohn’s disease involving the ileum. ISME J, 1(5), 403–418. doi:10.1038/ismej.2007.52 18043660

Blanton, L. V., Charbonneau, M. R., Salih, T., Barratt, M. J., Venkatesh, S., Ilkaveya, O., … Gordon, J. I. (2016). Gut bacteria that prevent growth impairments transmitted by microbiota from malnourished children. Science, 351(6275). doi:10.1126/science.aad3311 26912898

Blattner, F. R., Plunkett, G., 3rd, Bloch, C. A., Perna, N. T., Burland, V., Riley, M., … Shao, Y. (1997). The complete genome sequence of Escherichia coli K-12. Science, 277(5331), 1453–1462. doi:10.1126/science.277.5331.1453 9278503

Browning, D. F., Wells, T. J., Franca, F. L., Morris, F. C., Sevastsyanovich, Y. R., Bryant, J. A., … Henderson, I. R. (2013). Laboratory adapted Escherichia coli K-12 becomes a pathogen of Caenorhabditis elegans upon restoration of O antigen biosynthesis. Mol Microbiol, 87(5), 939–950. doi:10.1111/mmi.12144 23350972

Buffington, S. A., Di Prisco, G. V., Auchtung, T. A., Ajami, N. J., Petrosino, J. F., & Costa-Mattioli, M. (2016). Microbial Reconstitution Reverses Maternal Diet-Induced Social and Synaptic Deficits in Offspring. Cell, 165(7), 1762–1775. doi:10.1016/j.cell.2016.06.001 27315483

Capone, F., Bonsignore, L. T., & Cirulli, F. (2005). Methods in the analysis of maternal behavior in the rodent. Curr Protoc Toxicol, Chapter 13, Unit13 19. doi:10.1002/0471140856.tx1309s26 23045113

Charbonneau, M. R., O’Donnell, D., Blanton, L. V., Totten, S. M., Davis, J. C., Barratt, M. J., … Gordon, J. I. (2016). Sialylated Milk Oligosaccharides Promote Microbiota-Dependent Growth in Models of Infant Undernutrition. Cell, 164(5), 859–871. doi:10.1016/j.cell.2016.01.024 26898329

De Lucia Rolfe, E., de Franca, G. V. A., Vianna, C. A., Gigante, D. P., Miranda, J. J., Yudkin, J. S., … Ong, K. K. (2018). Associations of stunting in early childhood with cardiometabolic risk factors in adulthood. PLoS One, 13(4), e0192196. doi:10.1371/journal.pone.0192196 29641597

Ferdous, M., Zhou, K., Mellmann, A., Morabito, S., Croughs, P. D., de Boer, R. F., … Friedrich, A. W. (2015). Is Shiga Toxin-Negative Escherichia coli O157:H7 Enteropathogenic or Enterohemorrhagic Escherichia coli? Comprehensive Molecular Analysis Using Whole-Genome Sequencing. J Clin Microbiol, 53(11), 3530–3538. doi:10.1128/JCM.01899-15 26311863

Fernandez, C. A., Des Rosiers, C., Previs, S. F., David, F., & Brunengraber, H. (1996). Correction of 13C mass isotopomer distributions for natural stable isotope abundance. J Mass Spectrom, 31(3), 255–262. doi:10.1002/(SICI)1096-9888(199603)31:3<255::AID-JMS290>3.0.CO;2-38799277

Fischer, M., JeVenn, A., & Hipskind, P. (2015). Evaluation of muscle and fat loss as diagnostic criteria for malnutrition. Nutr Clin Pract, 30(2), 239–248. doi:10.1177/0884533615573053 25753808

Fuhrer, A., Sprenger, N., Kurakevich, E., Borsig, L., Chassard, C., & Hennet, T. (2010). Milk sialyllactose influences colitis in mice through selective intestinal bacterial colonization. J Exp Med, 207(13), 2843–2854. doi:10.1084/jem.20101098 21098096

Girma, S., Fikadu, T., & Abdisa, E. (2019). Maternal Common Mental Disorder as Predictors of Stunting among Children Aged 6-59 Months in Western Ethiopia: A Case-Control Study. Int J Pediatr, 2019, 4716482. doi:10.1155/2019/4716482 30956676

Gopalakrishna, K. P., Macadangdang, B. R., Rogers, M. B., Tometich, J. T., Firek, B. A., Baker, R., … Hand, T. W. (2019). Maternal IgA protects against the development of necrotizing enterocolitis in preterm infants. Nat Med, 25(7), 1110–1115. doi:10.1038/s41591-019-0480-9 31209335

Hawkes, C. P., & Grimberg, A. (2015). Insulin-Like Growth Factor-I is a Marker for the Nutritional State. Pediatr Endocrinol Rev, 13(2), 499–511. Retrieved from https://www.ncbi.nlm.nih.gov/pubmed/26841638 https://www.ncbi.nlm.nih.gov/pmc/articles/PMC5576178/pdf/nihms897988.pdf26841638

Hayashi, K., Morooka, N., Yamamoto, Y., Fujita, K., Isono, K., Choi, S., … Horiuchi, T. (2006). Highly accurate genome sequences of Escherichia coli K-12 strains MG1655 and W3110. Mol Syst Biol, 2, 2006 0007. doi:10.1038/msb4100049 16738553

Hintz, R. L., Suskind, R., Amatayakul, K., Thanangkul, O., & Olson, R. (1978). Plasma somatomedin and growth hormone values in children with protein-calorie malnutrition. J Pediatr, 92(1), 153–156. doi:10.1016/s0022-3476(78)80099-7 412936

Houghteling, P. D., & Walker, W. A. (2015). Why is initial bacterial colonization of the intestine important to infants’ and children’s health? J Pediatr Gastroenterol Nutr, 60(3), 294–307. doi:10.1097/MPG.000000000000059725313849

Hunt, K. M., Foster, J. A., Forney, L. J., Schutte, U. M., Beck, D. L., Abdo, Z., … McGuire, M. A. (2011). Characterization of the diversity and temporal stability of bacterial communities in human milk. PLoS One, 6(6), e21313. doi:10.1371/journal.pone.0021313 21695057

Kelleher, K. J., Casey, P. H., Bradley, R. H., Pope, S. K., Whiteside, L., Barrett, K. W., … Kirby, R. S. (1993). Risk factors and outcomes for failure to thrive in low birth weight preterm infants. Pediatrics, 91(5), 941–948. Retrieved from https://www.ncbi.nlm.nih.gov/pubmed/8474814 https://pediatrics.aappublications.org/content/pediatrics/91/5/941.full.pdf8474814

Kim, S., Kim, H., Yim, Y. S., Ha, S., Atarashi, K., Tan, T. G., … Huh, J. R. (2017). Maternal gut bacteria promote neurodevelopmental abnormalities in mouse offspring. Nature, 549(7673), 528–532. doi:10.1038/nature23910 28902840

Kloting, N., Koch, L., Wunderlich, T., Kern, M., Ruschke, K., Krone, W., … Bluher, M. (2008). Autocrine IGF-1 action in adipocytes controls systemic IGF-1 concentrations and growth. Diabetes, 57(8), 2074–2082. doi:10.2337/db07-1538 18443199

Koch, M. A., Reiner, G. L., Lugo, K. A., Kreuk, L. S., Stanbery, A. G., Ansaldo, E., … Barton, G. M. (2016). Maternal IgG and IgA Antibodies Dampen Mucosal T Helper Cell Responses in Early Life. Cell, 165(4), 827–841. doi:10.1016/j.cell.2016.04.055 27153495

Lee, W. N., Bassilian, S., Ajie, H. O., Schoeller, D. A., Edmond, J., Bergner, E. A., & Byerley, L. O. (1994). In vivo measurement of fatty acids and cholesterol synthesis using D2O and mass isotopomer analysis. Am J Physiol, 266(5 Pt 1), E699–708. doi:10.1152/ajpendo.1994.266.5.E699 8203508

Lee, W. N., Bassilian, S., Lim, S., & Boros, L. G. (2000). Loss of regulation of lipogenesis in the Zucker diabetic (ZDF) rat. Am J Physiol Endocrinol Metab, 279(2), E425–432. doi:10.1152/ajpendo.2000.279.2.E425 10913044

Ley, R. E., Backhed, F., Turnbaugh, P., Lozupone, C. A., Knight, R. D., & Gordon, J. I. (2005). Obesity alters gut microbial ecology. Proc Natl Acad Sci U S A, 102(31), 11070–11075. doi:10.1073/pnas.0504978102 16033867

Lindner, C., Thomsen, I., Wahl, B., Ugur, M., Sethi, M. K., Friedrichsen, M., … Pabst, O. (2015). Diversification of memory B cells drives the continuous adaptation of secretory antibodies to gut microbiota. Nat Immunol, 16(8), 880–888. doi:10.1038/ni.3213 26147688

Lukjancenko, O., Wassenaar, T. M., & Ussery, D. W. (2010). Comparison of 61 sequenced Escherichia coli genomes. Microb Ecol, 60(4), 708–720. doi:10.1007/s00248-010-9717-3 20623278

Moore, T., Beltran, L., Carbajal, S., Strom, S., Traag, J., Hursting, S. D., & DiGiovanni, J. (2008). Dietary energy balance modulates signaling through the Akt/mammalian target of rapamycin pathways in multiple epithelial tissues. Cancer Prev Res (Phila), 1(1), 65–76. doi:10.1158/1940-6207.CAPR-08-0022 19138937

Rahman, A., Lovel, H., Bunn, J., Iqbal, Z., & Harrington, R. (2004). Mothers’ mental health and infant growth: a case-control study from Rawalpindi, Pakistan. Child Care Health Dev, 30(1), 21–27. doi:10.1111/j.1365-2214.2004.00382.x 14678308

Reister, M., Hoffmeier, K., Krezdorn, N., Rotter, B., Liang, C., Rund, S., … Oelschlaeger, T. A. (2014). Complete genome sequence of the gram-negative probiotic Escherichia coli strain Nissle 1917. J Biotechnol, 187, 106–107. doi:10.1016/j.jbiotec.2014.07.442 25093936

Ross, J., Hanlon, C., Medhin, G., Alem, A., Tesfaye, F., Worku, B., … Prince, M. (2011). Perinatal mental distress and infant morbidity in Ethiopia: a cohort study. Arch Dis Child Fetal Neonatal Ed, 96(1), F59–64. doi:10.1136/adc.2010.183327 20667895

Schieber, A. M., Lee, Y. M., Chang, M. W., Leblanc, M., Collins, B., Downes, M., … Ayres, J. S. (2015). Disease tolerance mediated by microbiome E. coli involves inflammasome and IGF-1 signaling. Science, 350(6260), 558–563. doi:10.1126/science.aac6468 26516283

Schwarzer, M., Makki, K., Storelli, G., Machuca-Gayet, I., Srutkova, D., Hermanova, P., … Leulier, F. (2016). Lactobacillus plantarum strain maintains growth of infant mice during chronic undernutrition. Science, 351(6275), 854–857. doi:10.1126/science.aad8588 26912894

Shin, N. R., Whon, T. W., & Bae, J. W. (2015). Proteobacteria: microbial signature of dysbiosis in gut microbiota. Trends Biotechnol, 33(9), 496–503. doi:10.1016/j.tibtech.2015.06.011 26210164

Smith, H. W. (1975). Survival of orally administered E. coli K 12 in alimentary tract of man. Nature, 255(5508), 500–502. Retrieved from https://www.ncbi.nlm.nih.gov/pubmed/10942971094297

Smith, M. I., Yatsunenko, T., Manary, M. J., Trehan, I., Mkakosya, R., Cheng, J., … Gordon, J. I. (2013). Gut microbiomes of Malawian twin pairs discordant for kwashiorkor. Science, 339(6119), 548–554. doi:10.1126/science.1229000 23363771

Storelli, G., Defaye, A., Erkosar, B., Hols, P., Royet, J., & Leulier, F. (2011). Lactobacillus plantarum promotes Drosophila systemic growth by modulating hormonal signals through TOR-dependent nutrient sensing. Cell Metab, 14(3), 403–414. doi:10.1016/j.cmet.2011.07.012 21907145

Stratikopoulos, E., Szabolcs, M., Dragatsis, I., Klinakis, A., & Efstratiadis, A. (2008). The hormonal action of IGF1 in postnatal mouse growth. Proc Natl Acad Sci U S A, 105(49), 19378–19383. doi:10.1073/pnas.0809223105 19033454

Subramanian, S., Huq, S., Yatsunenko, T., Haque, R., Mahfuz, M., Alam, M. A., … Gordon, J. I. (2014). Persistent gut microbiota immaturity in malnourished Bangladeshi children. Nature, 510(7505), 417–421. doi:10.1038/nature13421 24896187

Surkan, P. J., Kennedy, C. E., Hurley, K. M., & Black, M. M. (2011). Maternal depression and early childhood growth in developing countries: systematic review and meta-analysis. Bull World Health Organ, 89(8), 608–615. doi:10.2471/BLT.11.088187 21836759

Suzawa, M., Muhammad, N. M., Joseph, B. S., & Bland, M. L. (2019). The Toll Signaling Pathway Targets the Insulin-like Peptide Dilp6 to Inhibit Growth in Drosophila. Cell Rep, 28(6), 1439–1446 e1435. doi:10.1016/j.celrep.2019.07.015 31390559

Ward, R. E., Ninonuevo, M., Mills, D. A., Lebrilla, C. B., & German, J. B. (2006). In vitro fermentation of breast milk oligosaccharides by Bifidobacterium infantis and Lactobacillus gasseri. Appl Environ Microbiol, 72(6), 4497–4499. doi:10.1128/AEM.02515-05 16751577

Weiss, G. A., & Hennet, T. (2012). The role of milk sialyllactose in intestinal bacterial colonization. Adv Nutr, 3(3), 483S–488S. doi:10.3945/an.111.001651 22585928

Xie, W., Jensen, S. K. G., Wade, M., Kumar, S., Westerlund, A., Kakon, S. H., … Nelson, C. A. (2019). Growth faltering is associated with altered brain functional connectivity and cognitive outcomes in urban Bangladeshi children exposed to early adversity. BMC Med, 17(1), 199. doi:10.1186/s12916-019-1431-5 31760950

Yan, J., Herzog, J. W., Tsang, K., Brennan, C. A., Bower, M. A., Garrett, W. S., … Charles, J. F. (2016). Gut microbiota induce IGF-1 and promote bone formation and growth. Proc Natl Acad Sci U S A, 113(47), E7554–E7563. doi:10.1073/pnas.1607235113 27821775

